# HDAC7 controls anti-viral and anti-tumor immunity by CD8^+^ T cells

**DOI:** 10.1101/2022.09.18.508452

**Authors:** Cansu Yerinde, Jacqueline Keye, Sibel Durlanik, Inka Freise, Franziska Nowak, Hsiang-Jung Hsiao, Marilena Letizia, Stephan Schlickeiser, Benedikt Obermayer, Adrian Huck, Marie Friedrich, Hao Wu, Désirée Kunkel, Anja A. Kühl, Sebastian Bauer, Andreas Thiel, Britta Siegmund, Rainer Glauben, Carl Weidinger

## Abstract

Class II histone deacetylases (HDAC) orchestrate T cell-dependent immune responses via the epigenetic control of genes and via the post-translational modification of cytoplasmic and nuclear proteins. However, the contribution of single HDAC family members to the differentiation and function of peripheral CD8^+^ T cells remains elusive. We here demonstrate that HDAC7-deficiency leads to the upregulation of immune checkpoint molecules, decreased calcium influx as well as increased apoptosis of peripheral murine CD8^+^ T cells, which we could link to a MEF2D-dependent induction of FasL expression ultimately deterring the survival of HDAC7-deficient CD8^+^ T cells. Likewise, we observed in mouse models of lymphoma, that mice with a T cell specific-deletion of *Hdac7* harbor impaired anti-tumor immune responses in syngeneic transfer models of lymphoma and we found that HDAC7 is required for CD8^+^ T cell-dependent memory recall responses in models of lymphocytic choriomeningitis virus infection. Taken together, we identify HDAC7 as a central regulator of cellular exhaustion and apoptosis of peripheral CD8^+^ T cells, controlling CD8^+^ T cell dependent anti-tumor and anti-viral immunity in mice.

**Significance:** Although HDAC7 was identified as an important regulator of thymocyte development and survival, its role in the homeostasis and the functions of adult CD8^+^ T cells is not fully understood. Here, we identify HDAC7 as a critical regulator of peripheral CD8^+^ T cells since its deletion impairs anti-tumor and anti-viral immune responses in mouse models of LCMV infection and transfer models of lymphoma. We attribute this phenotype to impaired survival, calcium homeostasis as well as deterred memory function and increased exhaustion of HDAC7-deficient CD8^+^ T cells. Our findings are of clinical relevance regarding potential immune suppressive side effects of HDAC inhibitors that are currently under clinical trials for the treatment of autoimmune diseases and cancers.

## Introduction

CD8^+^ T cells are essential for anti-tumor and anti-viral immunity by killing neoplastic and infected cells and by providing long-term immunity through the generation of memory T (T_mem_) cells. Upon antigen stimulation, naïve CD8^+^ T cells rapidly expand and differentiate into short-lived cytotoxic T cells (CTL), identified as killer cell lectin-like receptor subfamily G member 1 (KLRG1)^+^ and CD127^−^ cells^1^, which eradicate tumor target or virus-infected cells^2^. Following pathogen clearance, the population of KLRG1^+^CD127^−^CD8^+^ T cells contracts via undergoing apoptosis and leaves behind a small population of antigen-specific, long-lived KLRG1^−^CD127^+^ CD8^+^ memory T cells that can either rapidly give rise to new short-lived CTL, in case of antigen re-challenge^3^, or endure years until antigen re-exposure.

Several mechanisms have been described to play a role in the differentiation and maintenance of memory and effector cells e.g., the strength of antigen stimulation or the expression of specific transcription factors like Eomesodermin (Eomes) in CD8^+^ T_mem_ and T-bet in CTL^4,5^. Recent work has also highlighted the significant metabolic changes CD8^+^ T cells must undergo during the effector to memory cell transition^6^. Thus, high glycolytic activity favors the terminal differentiation and expansion of CTL, whereas decreased glycolytic flux enhances the formation of long-lived CD8^+^ T_mem_ cells^7^. First evidence also suggests that several amino acids and amino acid transporters control the differentiation and function of CD8^+^ T_mem_ cells via the regulation of the mammalian target of rapamycin (mTOR) signaling^8^. However, the pathways directing the metabolic switch in CD8^+^ T cells are not completely understood and it remains to be elucidated how these metabolic fate decisions are epigenetically controlled and imprinted in CD8^+^ T cells.

We have previously demonstrated that Store-operated Calcium Entry (SOCE) mediated by stromal interaction molecule (STIM)1, STIM2 as well as ORAI proteins, regulates effector functions of cytotoxic CD8^+^ T cells upon TCR activation such as the production of IFNγ, TNFα as well as the degranulation of toxic vesicles^9,10^. Additionally, SOCE is required for the differentiation, maintenance and re-activation of CD8^+^ T_mem_ cells during anti-viral response in lymphocytic choriomeningitis virus (LCMV) infected *Stim1*^*fl/fl*^*Stim2*^*fl/fl*^*Cd4-Cre* mice^11^. Importantly, Vaeth and colleagues recently showed that SOCE represents a central regulator in the metabolic switch of T cells, as SOCE-deficient T cells harbor a reduced activity of the Akt/mTOR pathway ultimately resulting in defective cellular expansion and differentiation upon antigen encounter^12^. It is currently not known how SOCE signaling components are regulated and which epigenetic and post-translational modifications influence its signaling strength in T cells.

Continuous antigen stimulation can drive CD8^+^ T cells into a dysfunctional and exhausted state, which stands in contrast to functional effector or memory cells generated upon the resolution of infections^13^. Thus, exhausted cells are incapable of clearing virus and cannot react with proper recall responses upon viral reinfection in mouse models of LCMV infections^13,14^. Besides chronic infections, T cell exhaustion was also found to play a central role in the immune evasion of tumors and still represents a major challenge in immune therapy of cancer^15^. The main characteristics of cellular exhaustion include the increased expression of multiple inhibitory checkpoint molecules such as PD-1, TIM-3, CTLA-4 and LAG-3 together with a decreased production of the effector cytokines IL-2, IFNγ and TNFα^16^. Moreover, exhausted CD8^+^ T cells possess distinct epigenetic, transcriptional and metabolic profiles suggesting that they might represent a distinct subset of effector and memory CD8^+^ T cells^16–19^. To date, the cellular signaling networks and epigenetic regulators of cellular exhaustion are incompletely understood. Over the last years, the modification of histone acetylation patterns has been described in several studies as a key regulator of T cell differentiation and function^20–23^. Among class II histone deacetylases (HDACs), HDAC7 has emerged as a prominent candidate that might not only control thymic development of T cells via the transcriptional regulation of apoptosis in thymic T cells^20,24^, but which might also be pivotal for the function of adult peripheral T cells^25^. Thereby, phosphorylation of HDAC7 and its subsequent nuclear export is observed upon TCR activation^26^. Due to its function as nuclear and cytoplasmic deacetylase, HDAC7 can play dual roles, both as a transcriptional repressor as well as a regulator of protein-protein interactions via post-translational modifications of cytoplasmic target proteins and nuclear transcription factors such as the myocyte enhancer factor 2 (MEF2)^25–31^. However, the functional role of HDAC7 in CD8^+^ T cell-mediated immune responses is so far unknown.

Here, we discovered that *Hdac7*-deficient CD8^+^ T cells display impaired SOCE as well as a deregulation of metabolism, exhaustion and apoptosis regulating genes, ultimately deterring the metabolic fitness and survival of CD8^+^ T cells *in vitro*. By using conditional *Hdac7* knockout mice, we could furthermore demonstrate that the lack of HDAC7 results in dysfunctional CD8^+^ T cell-dependent anti-tumor immune responses due to a decreased survival of tumor-infiltrating CD8^+^ T cells paralleled by an increased cellular exhaustion of *Hdac7*-deficient CD8^+^ T cells *in vivo*. Moreover, we observed that HDAC7 is required for the formation of virus-specific memory CD8^+^ T cells and proper recall responses in LCMV models of infection, as *Hdac7-*deficient virus-specific CD8^+^ T cells featured an enrichment of exhaustion-related transcriptional signatures and failed to properly expand upon viral re-challenge models ultimately leading to uncontrolled viral replication in mice. Taken together, we here provide evidence that HDAC7 controls the cellular exhaustion and the survival of peripheral CD8^+^ T cells and that HDAC7 is required for proper anti-tumor and anti-viral immune responses.

## Results

### Deletion of *Hdac7* in T cells results in a pre-activated phenotype of CD8^+^ T cells and does not affect the function and differentiation of the CD4^+^ T cell compartment

The epigenetic regulators controlling the homeostasis of adult CD8^+^ T cells are incompletely understood and the specific role of class II HDACs in the differentiation and function of effector and memory CD8^+^ T cells as well as their contribution to the regulation of cellular exhaustion remain elusive. To delineate which class II HDAC members are expressed in peripheral CD8^+^ T cells, we first determined the mRNA expression levels of class II HDAC members during CTL differentiation of wild type (Wt) CD8^+^ T cells *in vitro*. We observed that *Hdac7* was the most highly transcribed class II member at all investigated time points (**Fig. 1A**). To elucidate the role of HDAC7 in the differentiation and function of T cells, we subsequently generated conditional *Hdac7* knockout (*Hdac7*^*ko*^) mice by crossing *Hdac7*^*fl/fl*^ and *Cd4-Cre* mice, which resulted in a deletion of *Hdac7* both in CD4^+^ and CD8^+^ T cells (**Suppl. Fig. 1A**) and which did not cause any spontaneous phenotype in *Hdac7*^*ko*^ mice under steady-state conditions (data not shown). We next compared the splenic T cell composition of *Hdac7*^*ko*^ or Wt mice by mass cytometry, using two independent panels of 31 and 35 lineage and functional markers (**Suppl. Tab. 1, 2**) and by subsequently applying the t-SNE algorithm on pre-gated CD3^+^ T cells. Thereby, we identified a lack of naïve CD62L^+^CD44^−^CD8^+^ T cells in *Hdac7*^*ko*^ mice (**Suppl. Fig. 1B**). These phenotypical differences became even more evident, when the clustering of earth mover’s distance (EMD) values of all mass cytometry markers after *ex vivo* stimulation with PMA/ionomycin was analyzed. Here, CD8^+^ T cells of *Hdac7*^*ko*^ mice varied significantly from those of Wt mice (**Fig. 1B**), whereas CD4^+^ T cells displayed a closer similarity between both groups (**Suppl. Fig. 1C**), suggesting that HDAC7-deficiency is mainly affecting peripheral CD8^+^ T cells but not CD4^+^ T cells. Accordingly, Wt and *Hdac7*^*ko*^ CD4^+^ T cells shared the same colitogenic potential in an adoptive T cell transfer model of colitis, as no significant differences in disease severity could be detected between recipient *Rag2*^*-/*-^ mice that had either received naïve Wt or *Hdac7*^*ko*^ CD4^+^ T cells (**Suppl. Fig. 1D-G**).

**Figure 1.**
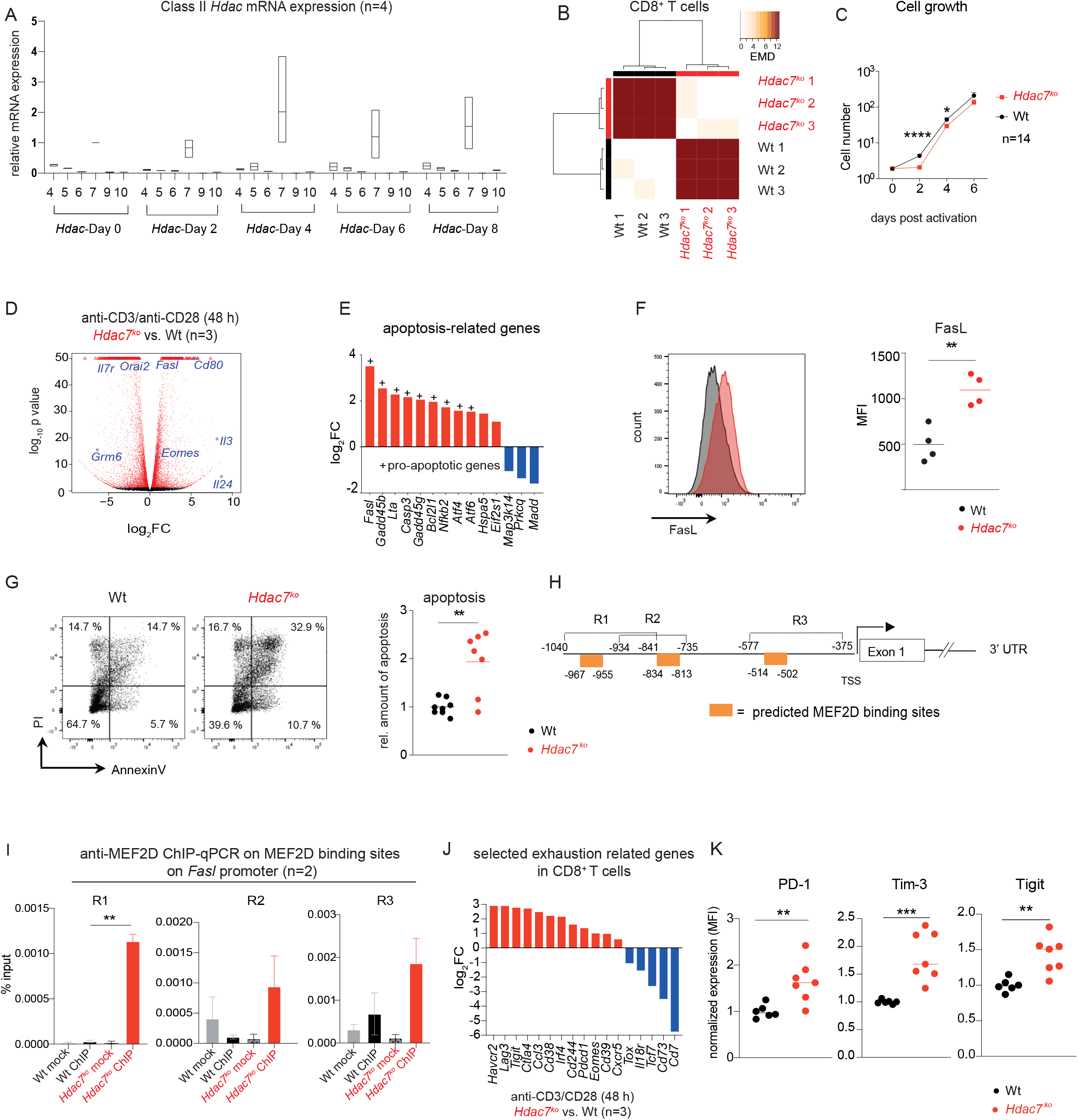
HDAC7 is crucial for the survival and the suppression of exhaustion markers in CD8^+^ T cells. **(A)** Box plots showing the mRNA expressions of Class II HDACs during *in vitro* CTL differentiation. Ct values were normalized to the Ct values of *36b4* and 2^−ΔCt^ was calculated. Expression of Class II HDACs are shown relative to the expression of *Hdac7* on Day 0 (n=4). **(B)** Heatmap displaying the pairwise earth-mover’s distance (EMD) values of the cellular density distribution within CD8^+^ T cell population isolated from the spleens of Wt littermates or *Hdac7*^*fl/fl*^*-Cd4-Cre*^*+*^*(Hdac7*^*ko*^) mice after PMA/ionomycin stimulation over a 2D t-SNE space (n=3 for both groups). **(C)** Cell growth curve after *in vitro* activation with anti-CD3/CD28 antibodies of CD8^+^ T cells from Wt or *Hdac7*^*ko*^ mice (n=14). **(D)** Volcano plots showing differentially regulated genes compared to Wt from comparative RNA-sequencing analysis of anti-CD3/CD28 antibody activated CD8^+^ T cells (n=3). **(E)** Fold-change expression of pro-apoptotic genes in *Hdac7*^*ko*^ CD8^+^ T cells in comparison to *in vitro* activated Wt CTL as assessed by RNA-sequencing (for all p<0.05, n=3). **(F)** Representative histograms showing FasL expression anti-CD3/CD28 stimulated Wt and *Hdac7*^*ko*^ CD8^+^ T cells 2 days post activation (left) and MFI values of activated Wt and *Hdac7*^*ko*^ CD8^+^ T cells (right; n=4, multiple t test). **(G)** Representative plots showing Annexin V and PI staining and the relative amount of apoptosis in Wt and *Hdac7*^*KO*^ CTLs (right) 7 days post activation. Plots are representative of 3 independent experiments. Total frequencies of Annexin V^+^PI^+^ cells were normalized to the mean value of Wt CTLs (n=7-8, multiple t test). **(H)** Schematics of putative MEF2D binding sites at the murine *Fasl* promoter identified by *in silico* analysis. **(I)** Chromatin immunoprecipitation (ChIP)-qPCR analysis of MEF2D binding on the *Fasl* promoter on the MEF2D binding sites R1-R3. CD8^+^ T cells from *Hdac7*^*ko*^ and Wt littermates were *in vitro* activated for 48 h with anti-CD3/CD28 antibodies and murine IL-2. ChIP was performed using anti-MEF2D antibody, followed by qPCR analysis. Plots show results of two independent experiments, in which 2-3 mice per group were pooled (multiple t test). **(J)** Fold change expression of CD8^+^ T cell exhaustion-related genes from comparative RNA-sequencing in *Hdac7*^*ko*^ vs. Wt CD8^+^ T cells 48 h after *in vitro* activation with anti-CD3/CD28 antibodies. (For all p<0.05, n=3) and **(K)** Dot plots showing normalized protein expression of PD-1, Tim-3 and Tigit in Wt and *Hdac7*^*ko*^ CD8^+^ T cells 48 h after *in vitro* activation with anti-CD3/CD28 antibodies. MFI values were normalized to the mean MFI of Wt. (3 independent experiments, n=6-7, multiple t-test) *p < 0.05; **p < 0.01, ***p < 0.001.

### Increased apoptosis in HDAC7-deficient CD8^+^ T cells

To better characterize the impact of HDAC7-deficiency on the homeostasis of peripheral CD8^+^ T cells, we next compared the expansion and the transcriptome of *Hdac7*^*ko*^ and Wt CD8^+^ T cells after *in vitro* activation with anti-CD3/CD28 antibodies. Interestingly, the cell number of *Hdac7*^*ko*^ CD8^+^ T cells was significantly reduced 2-4 days after initial stimulation (**Fig. 1C**) despite a comparable proliferative capacity of *Hdac7*-deficient and Wt CD8^+^ T cells (**Suppl. Fig. 2A-C**). RNA sequencing of *in vitro* activated naive CD8^+^ T cells from either Wt or *Hdac7*^*ko*^ mice revealed that several pro-apoptotic genes were upregulated in *Hdac7*^*ko*^ CD8^+^ T cells compared to Wt cells (**Fig. 1D/E**) of which FasL expression could also be validated on the protein level (**Fig. 1F**), ultimately leading to an increased rate of apoptosis and impaired survival of *Hdac7*^*ko*^ CD8^+^ T cells (**Fig. 1G**). Since the transcription factor MEF2D has been described to be directly regulated by HDAC7 and MEF2D is a known inducer of pro-apoptotic genes in T cells^20^, we hypothesized that the observed increased expression of FasL in *Hdac7*^*ko*^ CD8^+^ T cells might be due to an enhanced binding of MEF2D to the *Fasl* promoter. Likewise, we observed that MEF2D is significantly enriched in *Hdac7*^*ko*^ CD8^+^ T cells on three predicted binding sites (R1-R3) within the *FasL* promoter (**Fig. 1H/I**).

### HDAC7-dependent control of calcium homeostasis and metabolism

RNA sequencing analyses of *in vitro* synchronized CD8^+^ T cells from either Wt or *Hdac7*^*ko*^ mice furthermore revealed the transcriptional deregulation of several key components of SOCE-signaling in HDAC7-deficient CD8^+^ T cells including the significant downregulation of the calcium channel protein *Orai2*, the adaptor molecule *Stim1*, the sarco/endoplasmic reticulum Ca^2+^-ATPase (SERCA) pump *Atp2a2* as well as a down-regulation of the inositol-3-phosphate receptors (*Itpr*)*2* and *Itpr3*, highlighting that HDAC7 is required for proper Ca^2+^ homeostasis in CD8^+^ T cells as HDAC7-deficient CD8^+^ T cells also display a significantly decreased calcium influx after stimulation with thapsigargin when compared to Wt cells (**Suppl. Fig. 2D/E**).

Consistent with our RNA-seq data showing the upregulation of amino acid metabolism-related genes in *Hdac7*^*ko*^ CD8^+^ T cells, (**Suppl. Fig. 2F-H**), metabolic flux analyses revealed that while HDAC7-deficiency does not affect glycolysis and mitochondrial metabolism in CTLs, it results in increased glutamine uptake in HDAC7-deficient CTLs when compared to Wt CTLs (**Suppl. Fig. 2I-K**). In summary, our data demonstrate that HDAC7 has crucial roles for the survival, calcium homeostasis as well as the metabolic fitness of peripheral adult CD8^+^ T cells.

### HDAC7 is needed to suppress the upregulation of exhaustion markers during repeated antigen stimulation

Interestingly, we also observed a significant transcriptional upregulation of various well known CD8^+^ T cell exhaustion markers including *Havcr2, Ctla4, Pdcd1, Lag3* and *Tigit* by RNA sequencing in CD8^+^ T cells from *Hdac7*^*ko*^ mice 48 h after *ex vivo* stimulation with anti-CD3/CD28 antibodies, suggesting a crucial role for HDAC7 for the suppression of exhaustion markers in these cells (**Fig. 1J/K**).

To further investigate if HDAC7 contributes to the suppression of T cell exhaustion in CD8^+^ T cells, we next investigated the correlation of *Hdac7* expression with the expression of exhaustion markers in Wt CD8^+^ T cells in an *in vitro* model of chronic T cell stimulation^32^. Thus, we isolated CD8^+^ T cells from Wt mice and activated them for 48 hours with anti-CD3/CD28 antibodies in the presence of recombinant IL-2. On day 2, cells were divided into two groups: While the experimental group was exposed to repeated stimulation with anti-CD3/CD28 every other day to mimic chronic antigen stimulation, cells in the control group were grown in the presence of IL-2 only^32^ (**Fig. 2A** and **Suppl. Fig. 3A**). We first determined whether the expression of *Hdac7* and the other class II *Hdac* genes were affected by repeated stimulation in CD8^+^ T cells. Interestingly, the expression of *Hdac7* significantly decreased upon repeated re-stimulation of cells (**Fig. 2B**). Next, we analyzed how *Hdac7* expression was correlated with the expression of exhaustion marker genes. Significant negative correlations were observed between *Hdac7* expression and *Pdcd1, Havcr2* and *Tigit* (**Fig. 2C-E**), which were upregulated in Wt CD8^+^ T cells upon repeated anti-CD3/CD28 stimulation (**Suppl. Fig. 3B**). In accordance, we detected a significantly higher protein expressions of PD-1, Tim-3 and Tigit in *Hdac7*^*ko*^ CD8^+^ T cells compared to Wt when we applied our *in vitro* model of repeated T cell stimulation on *Hdac7*^*ko*^ CD8^+^ T cells (**Fig. 2F-H**). These results suggest that HDAC7 acts as a repressor of CD8^+^ T cell exhaustion-markers under chronic and repeated antigen stimulations.

**Figure 2.**
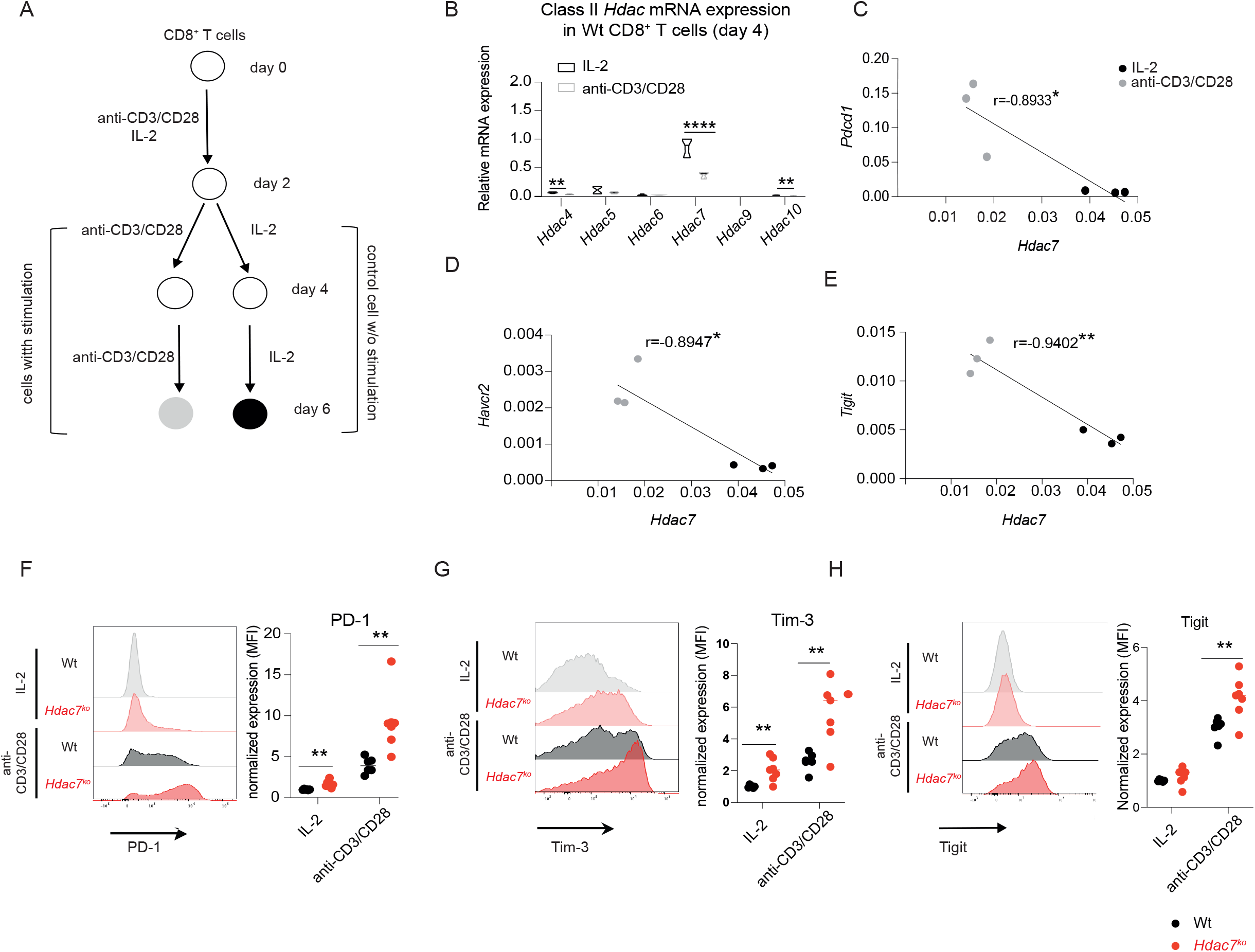
HDAC7 deletion in CD8^+^ T cells results in the upregulation of exhaustion markers during repeated antigen stimulation. **(A)** Experimental setup for *in vitro* chronic T cell stimulation model. Wt CD8^+^ T cells were *in vitro* activated with anti-CD3/CD28 antibodies for 48 h and cultured in the presence of recombinant IL-2 only or restimulated with anti-CD3/CD28 antibodies for an additional 48 h. **(B)** Relative mRNA expression of class II HDACs upon restimulation with anti-CD3/CD28 on day 4 relative to control cells grown in the presence of IL-2 only without repeated stimulation. Ct values were normalized to the Ct values of *36b4* and 2^−ΔCt^ was calculated (n=4, multiple t test). **(C-E)** Representative graphs showing the correlation between mRNA expressions of *Hdac7* and *Pdcd1, Havcr2* and *Tigit*. Pearson correlation analysis with 95% confidence interval (n=6). *p< 0.05; **p < 0.01, ***p< 0.001. **(F-H)** Representative primary FACS plots and dot plots showing the expression of **(F)** PD-1, **(G)** Tim-3 and **(H)** TIGIT in Wt or *Hdac7*^*ko*^ CD8^+^ T cells with or without restimulation *in vitro* (multiple t test, 3 independent experiments, n=6-7). **\***p<0.05, **p < 0.01, ***p < 0.001.

Remarkably, this increase in the expression of cellular exhaustion markers was also associated with increased apoptosis in CD8^+^ T cells lacking HDAC7 demonstrating that *Hdac7*^*ko*^ CD8^+^ T cells are more prone to develop apoptosis during chronic stimulation, which is also a hallmark of exhausted CD8^+^ T cells^16^ (**Suppl. Fig. 3C**). Taken together these data suggest that HDAC7 is not only required for the survival of CD8^+^ T cells during continuous stimulation, but that it is also crucial to prevent cellular exhaustion of CD8^+^ T cells.

### Impaired anti-tumor immunity in *Hdac7*^*ko*^ mice

To address if HDAC7 also controls T cell survival and exhaustion during chronic and repetitive antigen-exposure *in vivo*, we next explored how the deletion of *Hdac7* affects anti-tumor immune responses of CD8^+^ T cells in mice. Thus, we challenged *Hdac7*^*ko*^ mice or Wt littermates with syngeneic murine EG.7-Ova lymphoma cells (**Fig. 3A**), which resulted in increased tumor volume and tumor weight in *Hdac7*^ko^ mice (**Fig. 3B/C**). These results were confirmed in *Hdac7*^*fl/fl*^*-E8I-Cre* mice (**Suppl. Fig. 4A-C**), in which HDAC7 is exclusively deleted in CD8^+^ T cells, supporting a CD8^+^ T cell intrinsic function of HDAC7 in proper anti-tumor immune responses. As shown in **Fig. 3D/E**, significantly fewer tumor infiltrating CD8^+^ T cells could be detected in tumors of *Hdac7*^*ko*^ mice, which was paralleled by an increased frequency of CD8^+^Tim-3^+^ cells suggesting that HDAC7 might be required for either the expansion, survival, exhaustion and/or the homing of tumor-specific CD8^+^ T cells.

**Figure 3.**
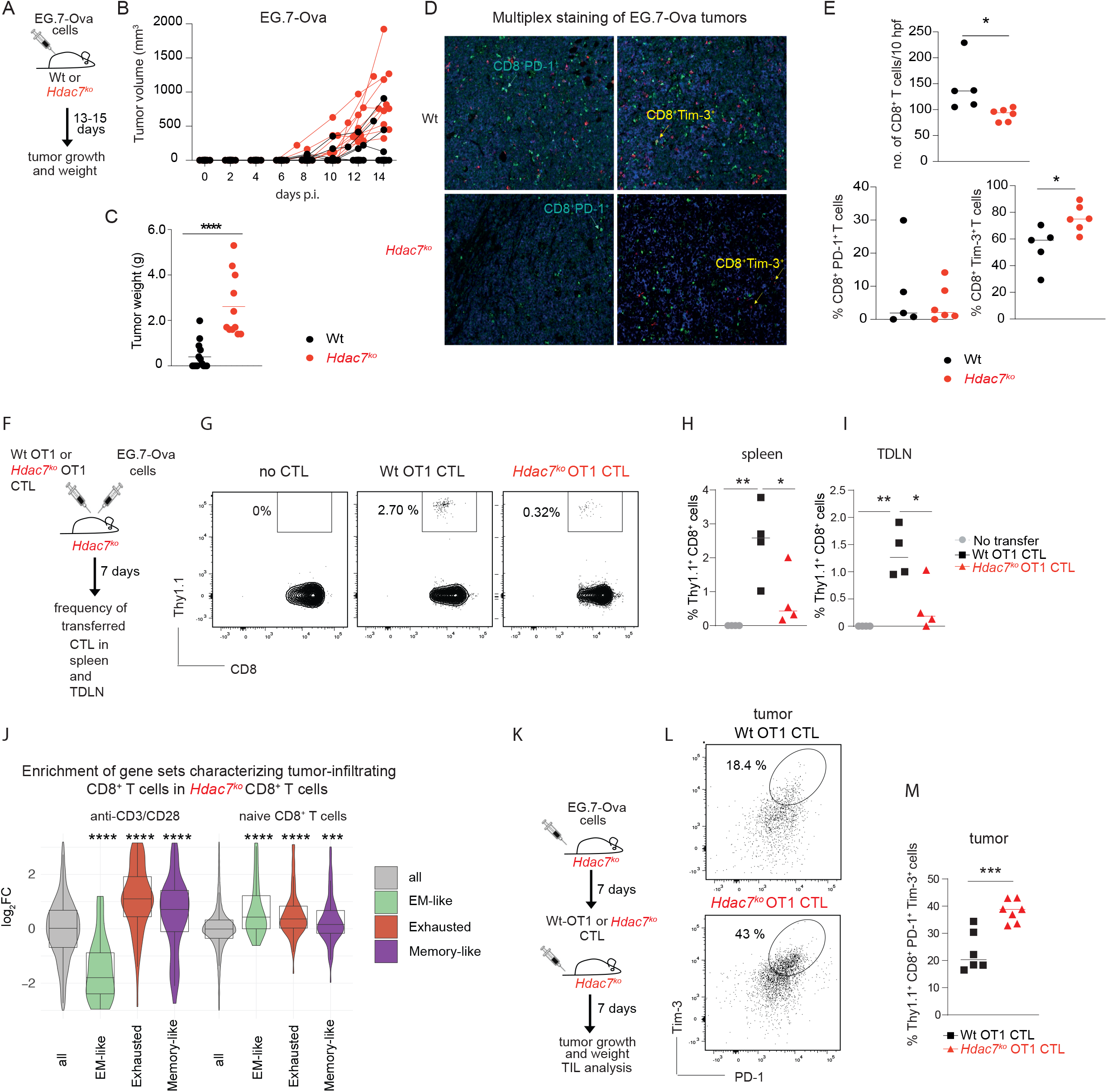
HDAC7 controls infiltration, *in vivo* persistence and the suppression of cellular exhaustion of CD8^+^ T cells in the tumors. **(A)** Wt and *Hdac7*^*fl/fl*^*Cd4-Cre (Hdac7*^*ko*^) mice were intradermally (i.d.) injected with 1×10^6^ EG.7-Ova cells. Tumor growth was followed for 14 days. **(B)** Tumor growth in Wt and *Hdac7*^*ko*^ mice shown as increase in tumor volumes (n=12-14 mice per group). **(C)** Tumor weight in Wt and *Hdac7*^*ko*^ mice on day 14 after tumor inoculation (n=12-14 animals per group, multiple t test). **(D)** Representative images from multiplex staining of EG.7-Ova tumor tissues in Wt (above) and *Hdac7*^*ko*^ (below) mice (CD8= green, PD-1=blue, Tim-3=yellow). **(E)** Number of tumor-infiltrating CD8^+^ T cells (upper panel) and the frequencies of CD8^+^PD-1^+^ and CD8^+^Tim-3^+^ cells (lower panels) per 5 high power fields (hpf) in EG.7-Ova tumors in Wt and *Hdac7*^*ko*^ mice 14 days p.i. (n=5-6 mice per group, multiple t test). **(F)** *Hdac7*^*ko*^ mice were injected with EG.7-Ova cells. Wt-OT1 or *Hdac7*^*ko*^-OT1 CTLs were transferred on the same day. The frequencies of transferred cells were analyzed after 7 days in spleens and tumor draining lymph nodes (TDLN). **(G)** Representative flow cytometry plots showing the splenic frequencies of Thy1.1^+^ Wt OT1 or Thy1.1^+^*Hdac7*^ko^-OT1 CTLs. (**H-I**) Dot plots of frequencies in spleens and TLDN of Wt OT1 and *Hdac7*^ko^-OT1 CTLs 7 days after i.v. transfer (n=4 mice per group, multiple t test). **(J)** *In silico* enrichment analysis of published gene sets^33^ obtained from tumor infiltrating CD8^+^ T cells in anti-CD3/C28 activated *Hdac7*^*ko*^ CD8^+^ T cells (Wilcoxon test). **(K)** *Hdac7*^*ko*^ mice were injected (i.d.) with 1×10^6^ EG.7-Ova cells. Seven days after tumor inoculation, recipient mice were injected with 7×10^6^ *in vitro* differentiated Wt OT1 or *Hdac7*^*ko*^ OT1 CTLs. Tumor growth was followed for 7 days post CTL transfer. **(L)** Representative FACS plots showing PD1^+^Tim3^+^Thy1.1^+^ CTLs in EG.7-Ova tumors in transferred mice. **(M)** Dot plot showing the frequencies of Thy1.1^+^PD1^+^Tim-3^+^CD8^+^ Wt OT1 and *Hdac7*^*ko*^ OT1 cells within EG.7-Ova tumors (n=6-7, multiple t test). *p< 0.05; **p < 0.01, ***p< 0.001, ****p<0.0001.

To exclude an impairment in the homing capacity of HDAC7-deficient CD8^+^ T cells, we transferred either *in vitro* expanded Wt *Thy1*.*1*^*+*^*Ot-1*^*+*^ (Wt-OT1) CTLs or *Hdac7*^*ko*^ *Thy1*.*1*^*+*^*Ot-1*^*+*^ (*Hdac7*^*ko*^-OT1) CTLs into recipient Wt mice with established tumors and compared the frequency of tumor-infiltrating CD8^+^ T cells 90 min after intravenous (i.v.) transfer (**Suppl. Fig. 4D**). As shown in **Suppl. Fig. 4E**, no statistically significant difference in the homing capacity to either tumors or tumor draining lymph nodes (TDLN) could be detected in mice receiving *Hdac7*^*ko*^ or Wt CTL. To clarify whether HDAC7 is required to maintain the survival of CD8^+^ T cells *in vivo*, we subsequently injected recipient *Hdac7*^*ko*^ mice with the same number of Thy1.1^+^ Wt-OT1 or *Hdac7*^*ko*^-OT1 CTLs and simultaneously challenged these mice with EG.7-Ova lymphoma cells (**Figure 3F**). Indeed, we observed a reduced frequency of transferred Thy1.1^+^ *Hdac7*^*ko*^ CTLs in spleens as well as TDLN, 7 days after adoptive transfer, implying that HDAC7 is needed for the *in vivo* persistence of transferred CTLs (**Figure 3G-I**). In accordance with our previous experiments, we also noted a significant enrichment of gene-signatures in the transcriptome of HDAC7-deficient CD8^+^ T cells that have been associated with the exhaustion of tumor-infiltrating CD8^+^ T cells^33^ (gene-signatures were derived from the re-analysis of published single-cell RNA sequencing data of melanoma infiltrating CD8^+^ T cells, **Fig. 3J** and **Suppl. Fig. 4F/G**). To confirm that the lack of HDAC7 in tumor infiltrating CTLs indeed results in an exhausted phenotype, CD8^+^ T cells from Wt OT1 or *Hdac7*^*ko*^ OT1 mice were *in vitro* differentiated into CTLs and subsequently i.v. injected into recipient EG.7-Ova tumor bearing *Hdac7*^*ko*^ mice (**Fig. 3K**), which significantly delayed tumor growth in both experimental groups when compared to EG7-Ova bearing mice not receiving CTLs (**Suppl. Fig. 4H/I**). However, the analysis of EG.7-Ova infiltrating CD8^+^ T cells revealed that tumor-infiltrating *Hdac7*^*ko*^ OT1 CD8^+^ T cells display a higher abundance of PD1^+^Tim3^+^ cells, implying that *Hdac7*-deficient CD8^+^ T cells are more prone to develop cellular exhaustion under chronic antigen stimulation within tumors (**Fig. 4L/M**), which could not be observed in transferred *Hdac7*^*ko*^ CD8^+^ T cells in spleens or TDLN (**Suppl. Fig. 4J/K**).

**Figure 4.**
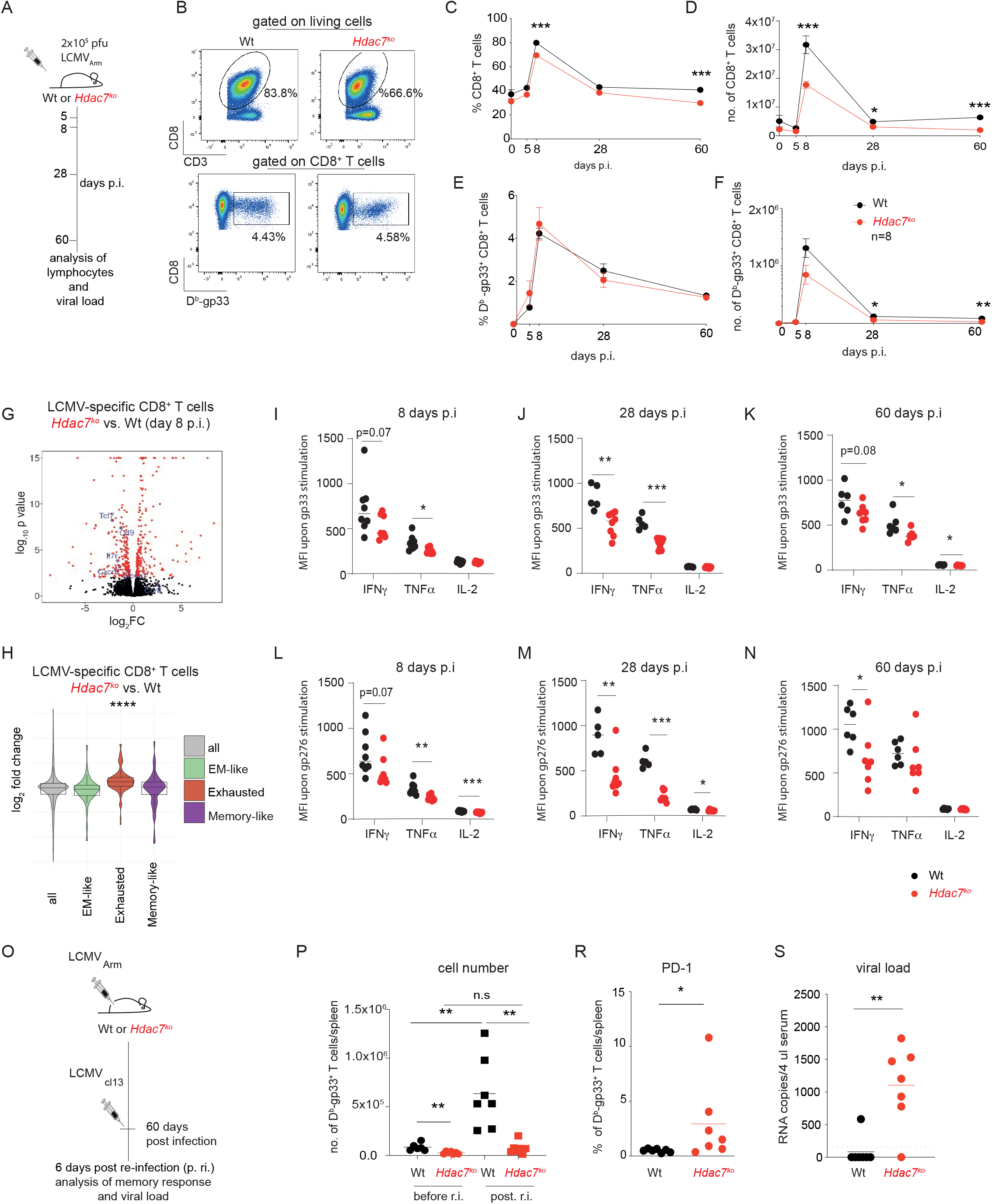
HDAC7 controls maintenance of CD8^+^ T cells and prevents cellular exhaustion during repeated LCMV infection. **(A)** Wt or *Hdac7*^*ko*^ mice were infected with 2×10^5^ plaque forming units (pfu) LCMV-Armstrong (LCMV_Arm_) virus by i.p. injections. Splenocytes from infected mice were analyzed by flow cytometry 5, 8, 28, and 60 days post infection (p.i.). **(B)** Representative plots gated on CD3^+^ T cells (upper panel) displaying the frequency of CD8^+^ T cells and D^b^-Gp33-streptamer+ CD3+CD8^+^ T cells (lower panel) of LCMV_Arm_ infected Wt and *Hdac7*^*ko*^ mice 8 days p.i.. **(C-F)** Frequencies (left) and total counts per spleen (right) of CD3^+^CD8^+^D^b^-Gp33-streptamer^+^ T cells at various time points p.i. (n=8, multiple t test). **(G)** Volcano plots showing differentially regulated genes compared to Wt from comparative RNA-sequencing analysis of LCMV-specific CD8^+^ T 8 days after LCMV infection (n=3). **(H)** *In silico* enrichment analysis of published gene sets^33^ obtained from tumor infiltrating CD8^+^ T cells in LCMV-specific CD8^+^ T cells 8 days after LCMV infection isolated from *Hdac7*^*ko*^ mice when compared to the respective Wt. **(I-K)** Mean fluorescence intensity (MFI) of IFNγ, TNFα, and IL-2 in Wt and *Hdac7*^*ko*^ CD8^+^ T cells after stimulation with D^b^-Gp33-peptide. **(L-N)** MFI of IFNγ, TNFα and IL-2 in Wt and *Hdac7*^*ko*^CD8^+^ T cells after stimulation with gp276-peptide on day 8, day 28 and day 60 p.i with LCMV_Arm_ (day 8 p.i.: n= 7-8; day 28 p.i.: n= 5-8; day 60 p.i.: 6-7 animals per group, multiple t test). **(O)** 60 days after primary infection with LCMV_Arm_, Wt or *Hdac7*^*ko*^ mice were re-challenged with LCMV-clone 13 (LCMVcl13). On day 6 post re-infection splenocytes were analyzed by flow cytometry. **(P)** Total counts per spleen of CD3^+^CD8^+^D^b^-Gp33-streptamer+ T cells in Wt and *Hdac7*^*ko*^ mice before and after LCMVcl13 re-infection (n=6-7, multiple t test). **(R)** Dot plots showing the frequency of PD-1^+^CD3^+^CD8^+^D^b^-Gp33-streptamer^+^ T cells in Wt and *Hdac7*^*ko*^ mice 6 days p.r.i. assessed by flow cytometry (n=7 per group, multiple t test). **(S)** Viral load in serum of LCMVcl13-infected Wt and *Hdac7*^*ko*^ mice after re-infection with LCMV_Arm_ (n=7, multiple *t*-test). *p < 0.05, **p<0.01, ***p<0.001

### Dysfunctional memory recall responses and increased cellular exhaustion in *Hdac7*^*ko*^ mice infected with LCMV

In case of chronic and repetitive antigen stimulation, CD8^+^ T cells progress into an exhausted phenotype at the expense of proper memory differentiation and responses^13,16^. Following up on the phenotypic differences of CD8^+^ T cells that we observed during our *in vivo* tumor experiments, we decided to functionally compare CD8^+^ T cell responses in *Hdac7*^*ko*^ and Wt mice over the course of acute and recurring viral infections with LCMV as a second model to understand the role of HDAC7 in CD8^+^ T cell-dependent immune responses against viruses. To this end, we first compared CD8^+^ T cell responses between *Hdac7*^*ko*^ mice and Wt littermates during acute infections with LCMV_Armstrong_ (LCMV_Arm_) (**Fig. 4A**). As shown in **Fig. 4B-D**, both the absolute number and the frequencies of total CD8^+^ T cells are significantly lower in *Hdac7*^*ko*^ mice compared to Wt littermates at all timepoints of infection including steady-state conditions. The frequencies of LCMV-specific D^b^-Gp33^+^CD8^+^ T cells were not changed in *Hdac7*^*ko*^ mice upon LCMV infection suggesting that the initial development of short lived D^b^-Gp33^+^ effector CD8^+^ T cells was unaffected by HDAC7-deficiency (**Fig. 4E**). However, in line with a general decrease in total CD8^+^ T cell numbers in *Hdac7*^*ko*^ mice, the absolute numbers of LCMV-specific D^b^-Gp33^+^CD8^+^ T cells were significantly lower in *Hdac7*^*ko*^ mice compared to Wt littermates 28 and 60 days post LCMV infection (**Fig. 4F**). Despite a lower absolute number of virus-specific CD8^+^ T cells, comparable copy numbers of LCMV_Arm_ could be detected in the serum of virus-infected *Hdac7*^*k*o^ mice and Wt littermates by quantitative (q)PCR (**Suppl. Fig. 5A**), highlighting that the initial effector functions of CTL are preserved in *Hdac7*^*ko*^ mice, despite an increased Eomes/Tbet ratio in *Hdac7*^*ko*^ mice (**Suppl. Fig. 5B-E**), suggesting a deterred transcriptional programming of *Hdac7*-deficient CD8^+^ T cells due to increased Eomes expression which has been described as a transcriptional marker of CD8^+^ T cell exhaustion^34,35,36^. Accordingly, comparative RNA-sequencing of LCMV-specific gp33^+^ CD8^+^ T cells isolated from LCMV-infected *Hdac7*^*ko*^ mice revealed an altered transcriptome compared to Wt littermates **(Figure 4G)**. When assessing the enrichment of published gene sets that have been implicated in the development and function of LCMV-specific CD8^+^ memory T cells^37^, we found that *Hdac7*-deficient, virus-specific CD8^+^ T cells displayed a significant enrichment of differentially expressed genes related to apoptosis, metabolism, transcription, signaling, effector functions, glycosylation, cell homing and the expression of cell surface receptors **(Suppl. Fig. 2F/G)**. Similar to *in vitro* activated *Hdac7*^*ko*^ CD8^+^ T cells and our observation in tumors, we furthermore observed a significant enrichment of published gene-signatures associated with the exhaustion of tumor-infiltrating CD8^+^ T cells in the transcriptome of LCMV-specific *Hdac7*-deficient CD8^+^ T cells^33^ (**Figure 4H**), further supporting that HDAC7 regulates cellular exhaustion of CD8^+^ T cells in anti-viral and anti-tumor immunity.

To analyze if the changes in cell differentiation observed in *Hdac7*-deficient CD8^+^ T cells during viral infections would also be reflected in altered cytokine patterns, we subsequently compared the production of IFNγ, TNFα and IL-2 in splenocytes of LCMV_Arm_-infected *Hdac7*^*ko*^ and Wt mice upon *ex vivo* stimulation with the immunodominant LCMV peptide Gp33 and the immune-subdominant Gp276 peptide^38,39^. In line with the pre-activated state of CD8^+^ T cells in *Hdac7*^*ko*^ mice under steady-state conditions, stimulation with Gp33 peptide induced a significantly higher frequency of IFNγ and TNFα producing cells in *Hdac7*-deficient CD8^+^ T cells 5 days p.i. but an equivalent fraction of IFNγ, TNFα and IL-2 producing cells 8 days p.i. (**Suppl. Fig. 6A-C**). Upon gp276-peptide stimulation, *Hdac7*^*ko*^ mice displayed decreased frequencies of all three cytokines for all timepoints we analyzed (**Suppl. Fig. 6D-F**). Moreover, in line with an exhaustion-like phenotype, the capacity to produce cytokines were severely impaired in *Hdac7*^*ko*^ CD8^+^ T cells upon *ex vivo* re-stimulation with both the gp33 and the gp276 peptides as HDAC7-deficient T cells displayed significantly reduced mean fluorescence intensities (MFI) of IFNy, TNFα and IL-2 (**Fig. 4I-N**), which is considered as a key feature of T cell exhaustion^40^. Additionally, a significant reduction in the total number of cytokine-producing cells was detected at almost all indicated time points, which can be explained by an additive effect of a lower frequency and absolute numbers of CD8^+^ T cells (**Suppl. Fig 6G-L**).

Until now, we had observed an increased expression of exhaustion markers in *Hdac7*^*ko*^ CD8^+^ T cells during repetitive stimulation *in vitro* as well as during chronic antigen exposure in tumors. To better understand how *Hdac7*^*ko*^ CD8^+^ T cells respond to repeated antigen exposure *in vivo*, we next analyzed the efficacy of *Hdac7*^*ko*^ mice to handle secondary infections after prior immunization. Thus, Wt or *Hdac7*^*ko*^ mice were infected with LCMV_Arm_ and subsequently challenged with LCMV clone 13 (LCMV_cl13_) 60 days after initial infection. Spleens of infected mice were then harvested 6 days post re-infection (p.ri.) with LCMV_cl13_ (**Fig. 4O**). *Hdac7*-deficient CD8^+^ T_mem_ cells failed to expand properly. While D^b^-Gp33^+^ CD8^+^ T Wt CD8^+^ T cells expanded 9 fold upon re-exposure to virus, *Hdac7*^*ko*^ CD8^+^ T cells expanded only 3-fold upon antigen re-encounter (**Fig. 4P**). Moreover, D^b^-Gp33^+^ *Hdac7*^*ko*^ cells displayed a significantly higher expression of the exhaustion marker PD-1 (**Fig. 4R**), which was paralleled by a significantly higher viral load 6 days post re-infection (**Fig. 4S**), further supporting that HDAC7 is required to suppress cellular exhaustion in CD8^+^ T cells and for maintaining CD8^+^ T cell dependent immune responses upon repetitive antigen-exposure during both anti-tumor and anti-viral immunity.

## Discussion

Taken together, we here identified HDAC7 as a crucial regulator of apoptosis, Ca^2+^ homeostasis, immune metabolism and cellular exhaustion, that is required for CD8^+^ T cell-dependent anti-viral and anti-tumor immunity. Thus, we not only observed that HDAC7 is critical for anti-tumor immunity against lymphoma cells but that it is furthermore pivotal for the proper development and maintenance of virus-specific memory CD8^+^ T cells and subsequently for effective memory recall responses.

Unexpectedly, we observed a pre-activated phenotype under steady state conditions in CD8^+^ T cells and a transcriptional upregulation of several exhaustion markers including Tim-3, PD-1, Tigit, Ctla-4, and Lag-3 upon stimulation of *Hdac7*^*ko*^ CD8^+^ T cells. Furthermore, *Hdac7*^*ko*^ mice featured a significantly reduced frequency of CD8^+^ T cells, which was paralleled by an increased amount of apoptosis in *Hdac7*-deficient CD8^+^ T cells upon activation *in vitro*. Accordingly, we observed a transcriptional deregulation of several pro-apoptotic genes in CD8^+^ T cells of *Hdac7*^*ko*^ mice, which was in line with previous findings in T cells expressing an inactive mutant of *Hdac7*. Here, in CD8^+^ T cells displayed a deregulation of apoptosis in the thymus and subsequently an impaired thymic selection of T cells^24^. Our current data suggest that HDAC7 controls the induction of apoptosis in peripheral CD8^+^ T cells in a FasL-dependent manner at least partially by regulating the binding of the transcription factor MEF2D to the *Fasl* promoter. We also observed increased apoptosis of *Hdac7*^*ko*^ CD8^+^ T cells upon repetitive stimulation *in vitro*, paralleled by an increased expression of key exhaustion markers including PD-1, Tim-3 and Tigit. Adoptive transfer experiments in tumor bearing mice also revealed that the lack of HDAC7 both impairs the persistence of CTLs *in vivo* as well as the suppression of PD-1 and Tim-3 expression by CTLs in the tumor microenvironment, which are the hallmarks of CD8^+^ T cell exhaustion, further implying that HDAC7 is a crucial factor to control the exhaustion and survival of CD8^+^ T cells.

Astonishingly, the CD4^+^ T cell compartment of LCMV-infected *Hdac7*^*ko*^ mice was largely unaffected and no significant differences in the colitogenic potential of HDAC7-deficient naive CD4^+^ T cells could be detected as assessed in a T-cell transfer model of colitis. Therefore, CD4^+^ T cells can sufficiently provide help to CD8^+^ T cells, suggesting that the observed defects in anti-viral immunity and anti-tumor immunity in *Hdac7*^*ko*^ mice are due to CD8^+^ T cell intrinsic functions of HDAC7. We also found that HDAC7 is also required for the function of memory CD8^+^ T cells. Thus, *Hdac7*-deficient CD8^+^ T cells revealed a defective cytokine production of TNFα, IFNγ and IL-2. Importantly, *Hdac7*-deficient memory T cells failed to give rise to effector CD8^+^ T cells upon re-infection with LCMV_cl13_, which resulted in a defective viral clearance and an induction of the cellular exhaustion marker PD-1 in virus-specific CD8^+^ T cells of *Hdac7*^*ko*^ mice. Furthermore, virus-specific *Hdac7*^*ko*^ CD8^+^ T cells displayed an impaired cellular expansion as well as a significantly decreased total number in *Hdac7*^*ko*^ mice upon re-infection with LCMV_cl13_. These observations are also in line with the exhaustion-like phenotype of *Hdac7*^*ko*^ CD8^+^ T cells during both *in vitro* stimulations and chronic antigen exposure within tumors in which *Hdac7*^*ko*^ CD8^+^ T cells cannot handle repetitive stimulations properly due to an increased expression of exhaustion markers and impaired *in vivo* persistence.

In line with these functional impairments, *Hdac7*-deficient CD8^+^ T cells were characterized by a reduced Ca^2+^ influx and a transcriptional deregulation of several SOCE-signaling components. The latter has been described to play a pivotal role not only in the regulation of T cell metabolism^12^ but also in the function and maintenance of CD8^+^ T cell-dependent memory responses^11^. Thus, *Stim1*^*fl/fl*^*Stim2*^*fl/f*^*CD4-Cre* mice display a similar phenotype as *Hdac7*^*ko*^ mice upon viral infection with LCMV as they feature exhausted CD8^+^ T cells with high PD-1 and Tim-3 expression as well as impaired production of IL-2, TNFα and IFNγ, despite an elevated Eomes expression^11^. Additionally, they are also incapable of mounting functional recall responses against reinfection with LCMV_cl13_ and show severe defects in anti-tumor immune responses in mouse models of melanoma and lymphoma^9^.

In regard of the critical function of HDAC7 for the survival of functional CD8^+^ T cells in mice and the observation that HDAC7 controls the expression of exhaustion markers including *Pdcd1, Havcr2, Tigit, Ctla4* and *Lag3* in murine CD8^+^ T cells, downregulation or functional inhibition of HDAC7 might represent an escape mechanism for human tumors and pathogens resulting in impaired anti-tumor and anti-viral immunity.

The dependence of anti-viral memory responses and anti-tumor immune functions on HDAC7 has critical implications for the clinical evaluation of pharmacologic HDAC inhibitors that are currently being tested for the treatment of autoimmune diseases and various types of cancer^41^. Thus, the pan-HDAC inhibitors vorinostat and panobinostat were shown to decrease IFNγ production, the survival and the cytotoxic capacity of human HIV-specific CD8^+^ T cells^42^. Thereby, HDAC inhibitors do not only block the enzymatic activity of HDAC7 but suppress the expression of HDAC7 as shown for various cell types including fibroblasts, epithelial cells, bladder and prostate cancer cells as well as myeloma cells^43^. Likewise, we observed a downregulation of HDAC7 after treatment of CD8^+^ T cells with vorinostat (**Suppl. Fig. 7A/C**), significantly reducing the protein expression of HDAC7 and triggering the induction of apoptosis^44^. Moreover, we observed an increase in the acetylation of H3 at lysine 9 (K9) and K14 residues at the global level in *Hdac7*^*ko*^ CD8^+^ T cells suggesting that HDAC7 might regulate all these processes through histone deacetylation (**Suppl. Fig. 7B**). In light of the growing number of HDAC inhibitors that are currently tested for the treatment of autoimmune and neoplastic diseases, our findings caution that inhibition of HDAC7 interferes with the development and maintenance of appropriate memory responses e. g., upon vaccination and might thus alter the function, the metabolic fitness and the survival of CD8^+^ T cells during immune responses against viral infections and tumors.

## Acknowledgement

We thank Erik Olsen for kindly providing Hdac7^fl/fl^ mice and Sergei Nedospasov for the generous gift of *Cd4-Cre* mice. Rag2^−/-^ mice were a gift from T. Blankenstein (MDC, Berlin). We thank Ahmed Hegazy and Daniel Herranz for the critical discussion of our data. We would like to acknowledge the technical assistance of Beate Kruse as well as Manuela Dingeldey, Pauline Piguet and Yasmina Rodriguez Sillke for assistance during experiments. The authors thank the Deutsche Forschungsgemeinschaft (DFG) for supporting the acquisition of the cell sorter that was used in the present study (INST 335/597-1).

## Funding

This work was funded by the Helmholtz Alliance (Preclinical Comprehensive Cancer Center to B. Siegmund), the DFG (We 5303/3-1 to C. Weidinger, TRR241 B01 to B. Siegmund and C. Weidinger, and SI 749/5-3 to B. Siegmund), the Thyssen-Foundation (to C. Weidinger) and the German-Israeli Foundation for Scientific Research and Development as well as the German Cancer Consortium (DKTK) to B. Siegmund and R. Glauben. J. Keye was funded for one year by the Sonnenfeld-Stiftung, C. Yerinde was supported by the Berlin School of Integrative Oncology (BSIO) and received funding by a Ruth Jeschke Scholarship for Tumor Immunology for one year. C. Weidinger received funding from the Berlin Institute of Health (BIH) Clinician Scientist Program.

## Data availability

All RNA sequencing data will be made publicly available upon acceptance.

## Disclosure

The authors declare no conflict of interest.

## Author Contributions

C. Y., J.K., S.D., I.F., F.S., H.H., M.L., M.F., H.W., D.K. and A.A.K. performed experiments, C.Y., J.K., S.D., I.F., S.S., S.B., A.T., B.S., R.G. and C.W. designed and analyzed experiments. C.Y., J.K., R.G. and C.W. wrote the paper.

## Online Materials and Methods

Detailed experimental methods including mass cytometry, tumor allografts, adoptive T cell transfer experiments, LCMV infections are available in Supplementary Information (SI) Appendix. All animal protocols were approved by the regional animal study committee of Berlin (LaGeSo, Berlin, Germany) and conducted accordingly.

## Notes

### Competing Interest Statement

The authors have declared no competing interest.

