## Supplementary data and methods for "HDAC7 controls anti-viral and anti-tumor immunity by CD8^+^ T cells"

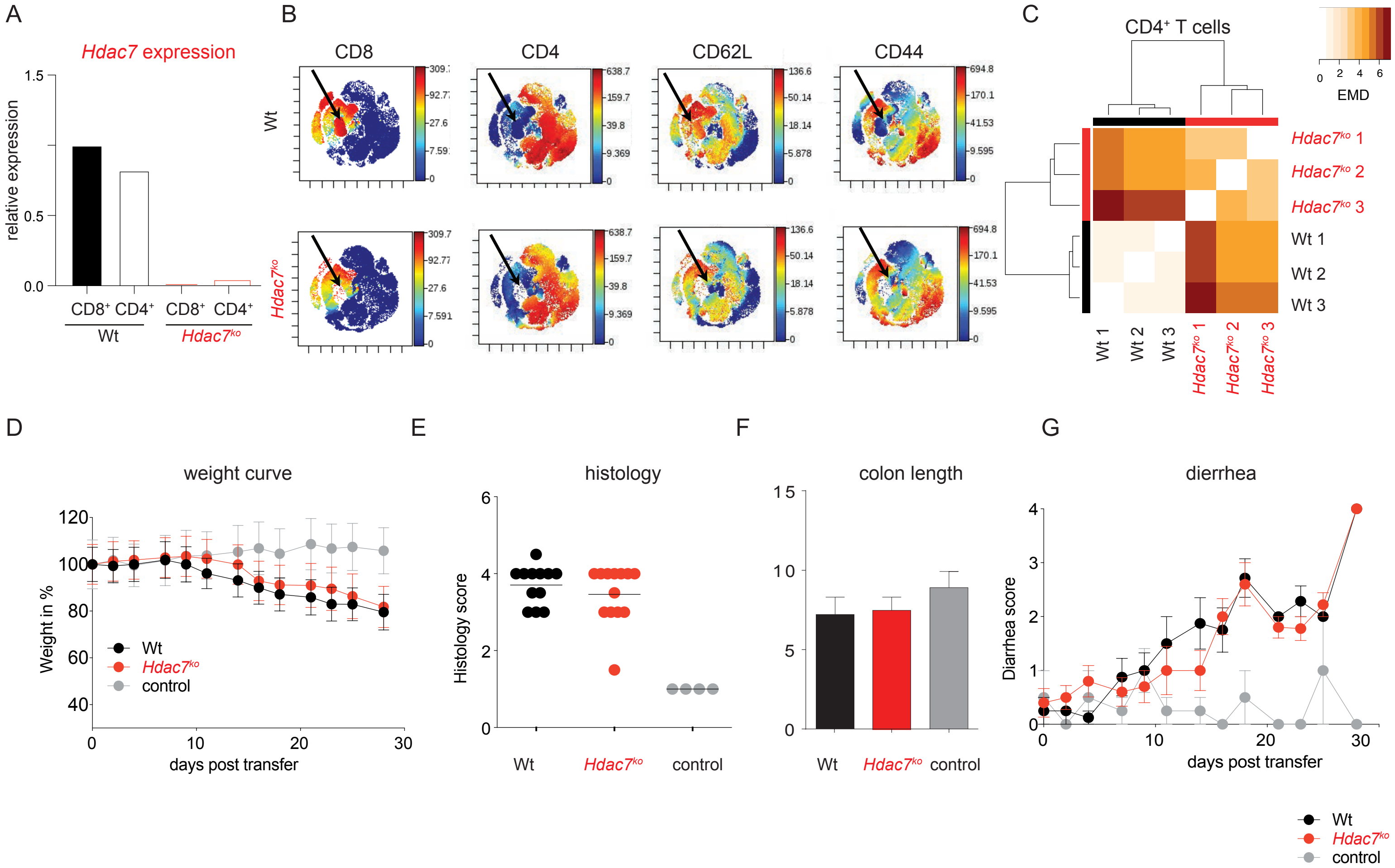


**Supplementary Figure 1: *Hdac7$^{fl/fl}$CD4-Cre* mice have a preactivated phenotype of CD8$^{+}$ T cells and an unaffected CD4$^{+}$ T cell compartment. (A)** Deletion efficiency of *Hdac7* in *Hdac7$^{fl/fl}$CD4-Cre* (*Hdac7$^{ko}$*) mice. CD4$^{+}$ and CD8$^{+}$ T cells were isolated from spleens of *Hdac7$^{ko}$* mice and Wt littermates. qPCR was performed to check the deletion efficiency of *Hdac7*. *Hdac7* expression was normalized to β-actin expression. Representative column plots of two independent experiments performed with two biologically independent samples per group. **(B)** Representative 2D-tSNE analyses of mass cytometry data comparing the distribution of CD8, CD4, CD62L and CD44 expressing cells in pre-gated CD45$^{+}$CD3$^{+}$ T cells isolated from the spleens of Wt littermates or *Hdac7$^{fl/fl}$-Cd4-Cre* (*Hdac7$^{ko}$*) mice (n=3 for both groups). **(C)** Heatmap displaying the pairwise earth-mover's distance (EMD) values of the cellular density distribution within CD4$^{+}$ T cell population after PMA/ionomycin stimulation over a 2D t-SNE space. **(D-G)** Transfer colitis experiments. 4x10$^{5}$ naïve CD4$^{+}$ T cells from Wt or *Hdac7$^{ko}$* mice were i.p. injected into *Rag2$^{-/-}$* mice to induce transfer colitis. Control *Rag2$^{-/-}$* mice were injected with PBS only. Mice were weighed and scored every second day and sacrificed on day 28 post transfer. **(D)** Line graphs showing the percentage of weight changes over the time of the disease. **(E)** Dot plots displaying the histology score composed of cell infiltration and tissue damage. **(F)** Bar graphs summarizing the colon length in cm. **(G)** Line graphs depicting the diarrhea score assessed by the consistency of feces.

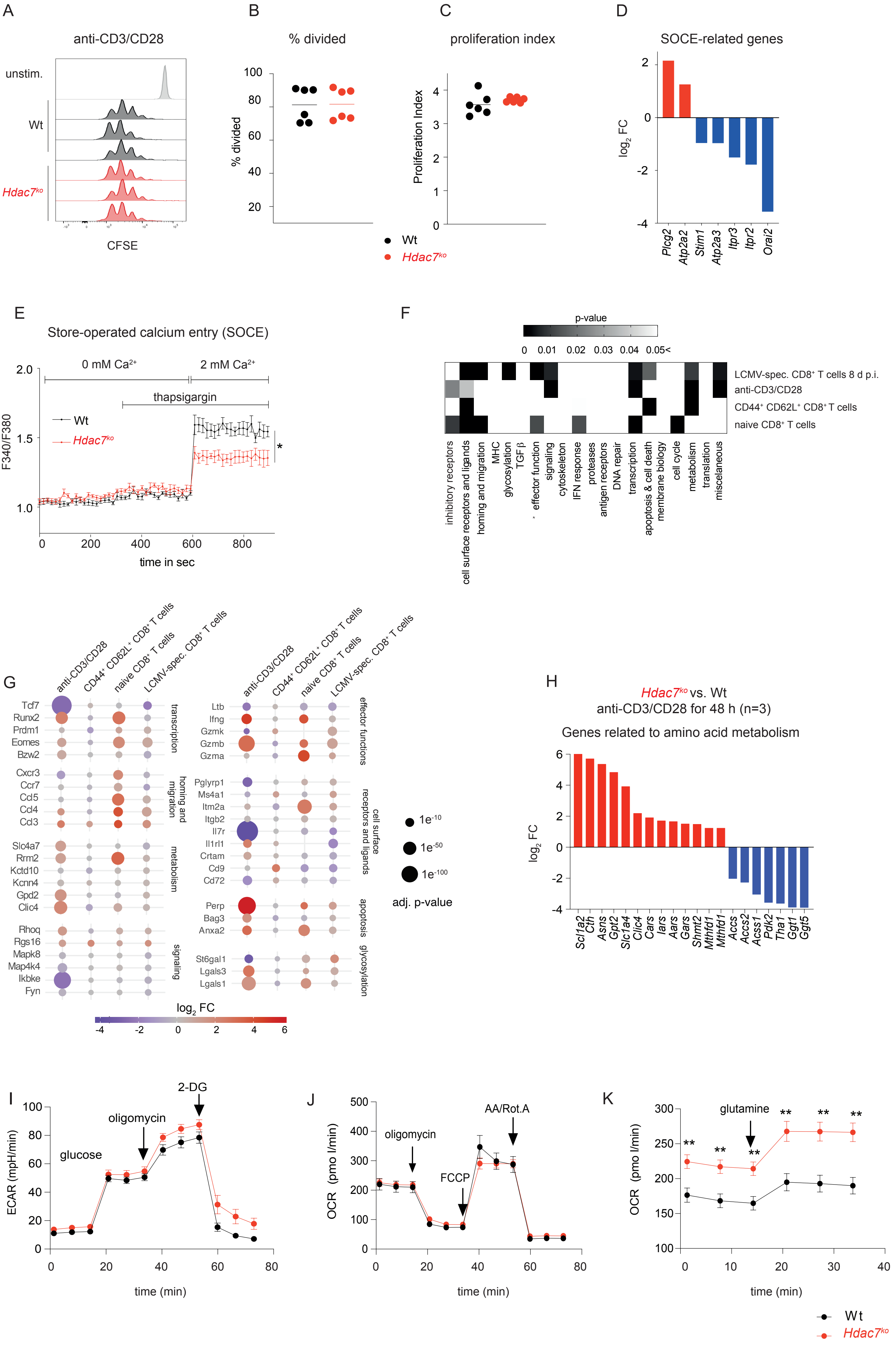
A
anti-CD3/CD28
unstim.
Wt
Hdac7ko
CFSE
B
% divided
% divided
C
proliferation index
Proliferation Index
Wt
Hdac7ko
D
SOCE-related genes
log2 FC
Plcg2
Atp2a2
Stim1
Atp2a3
Itpr3
Itpr2
Orai2
E
Store-operated calcium entry (SOCE)
0 mM Ca2+
2 mM Ca2+
thapsigargin
Wt
Hdac7ko
F340/F380
time in sec
F
p-value
0 0.01 0.02 0.03 0.04 0.05<
LCMV-spec. CD8+ T cells 8 d p.i.
anti-CD3/CD28
CD44+ CD62L+ CD8+ T cells
naive CD8+ T cells
inhibitory receptors
cell surface receptors and ligands
homing and migration
MHC
glycosylation
TGFβ
effector function
signaling
cytoskeleton
IFN response
proteases
antigen receptors
DNA repair
transcription
apoptosis & cell death
membrane biology
cell cycle
metabolism
translation
miscelaneous
G
anti-CD3/CD28
CD44+ CD62L+ CD8+ T cells
naive CD8+ T cells
LCMV-spec. CD8+ T cells
Tcf7
Runx2
Prdm1
Eomes
Bzw2
transcription
Cxcr3
Ccr7
Ccl5
Ccl4
Ccl3
homing and migration
Slc4a7
Rrm2
Kctd10
Kcnn4
Gpd2
Clic4
metabolism
Rhoq
Rgs16
Mapk8
Map4k4
Ikbke
Fyn
signaling
Ltb
Ifng
Gzmk
Gzmb
Gzma
effector functions
Pglyrp1
Ms4a1
Itm2a
Itgb2
Il7r
Il1rl1
Crtam
Cd9
Cd72
cell surface receptors and ligands
Perp
Bag3
Anxa2
apoptosis
St6gal1
Lgals3
Lgals1
glycosylation
1e-10
1e-50
1e-100
adj. p-value
log2 FC
H
Hdac7ko vs. Wt
anti-CD3/CD28 for 48 h (n=3)
Genes related to amino acid metabolism
log2 FC
Scl1a2
Cth
Asns
Gpt2
Slc1a4
Clic4
Cars
Iars
Aars
Gars
Shmt2
Mthfd1
Mthfd1l
Accs
Accs2
Acss1
Pdk2
Tha1
Ggt1
Ggt5
I
glucose
oligomycin
2-DG
ECAR (mpH/min)
time (min)
J
oligomycin
FCCP
AA/Rot.A
OCR (pmo l/min)
time (min)
K
glutamine
OCR (pmo l/min)
time (min)
Wt
Hdac7ko

**Supplementary Figure 2: HDAC7 deletion in $CD8^+$ T cells results in disturbed Store-Operated Calcium Entry (SOCE) and amino acid metabolism. (A)** Representative histograms of CFSE staining measured by flow cytometry 3 days post *in vitro* anti-CD3/CD28 activation of Wt and *Hdac7*$^{ko}$ $CD8^+$ T cells (two independent experiments, n=6). **(B)** Dot plots showing the percentage of divided cells, and **(C)** the proliferation index during CFSE dilution assay (n=6). Percentage of divided cells and the proliferation index were calculated by FlowJo software. **(D)** Fold-change expression of SOCE-component genes from comparative RNA-sequencing of Wt and *Hdac7*$^{ko}$ $CD8^+$ T cells activated with anti-CD3/CD28 antibodies for 48 h (for all $p<0.05$, n=3 per group). **(E)** Calcium influx in Wt and *Hdac7*$^{ko}$ CTLs, mean ± SEM of $Ca^{2+}$ influx rates (n=3 independent experiments, performed in duplicates, multiple t test). **(F)** Heatmaps showing the statistically significant enrichment of differentially regulated biological pathways in the different subsets of $CD8^+$ T cells between Wt or *Hdac7*$^{ko}$ mice, that have been previously implicated in the development and function of LCMV-specific $CD8^+$ memory T cells[7], and **(G)** top ten genes deregulated in these biological pathways in anti-CD3/CD28 activated, $CD44^+CD62L^+$, naïve and LCMV-specific $CD8^+$ *Hdac7*$^{ko}$ $CD8^+$ T cells. **(H)** Fold change expression of amino acid metabolism related genes in *Hdac7*$^{ko}$ $CD8^+$ T cells compared to Wt cells from RNA-sequencing (for all $p<0.05$, n=3 per group). **(I-K)** Metabolic flux measurements of Wt and *Hdac7*$^{ko}$ CTLs 7 days post activation. **(J)** Glycolysis levels as assessed by extracellular acidification rate (ECAR) values measured in response to 10 mM glucose, 1 µM oligomycin and 50 mM 2-DG treatment (n=3, biologically independent samples). **(K)** Mitochondrial respiration levels assessed by oxygen consumption rate (OCR) values measured in response to 2 µM oligomycin, 1 M FCCP and 0.5 M RotA/AA treatment (n=3, biologically independent samples). **(L)** Glutamine uptake assessed by OCR levels measured in response to 4 mM glutamine treatment. Error bars indicate mean ± SEM. (n=3, two-way Anova test) $^{**}p<0.01$

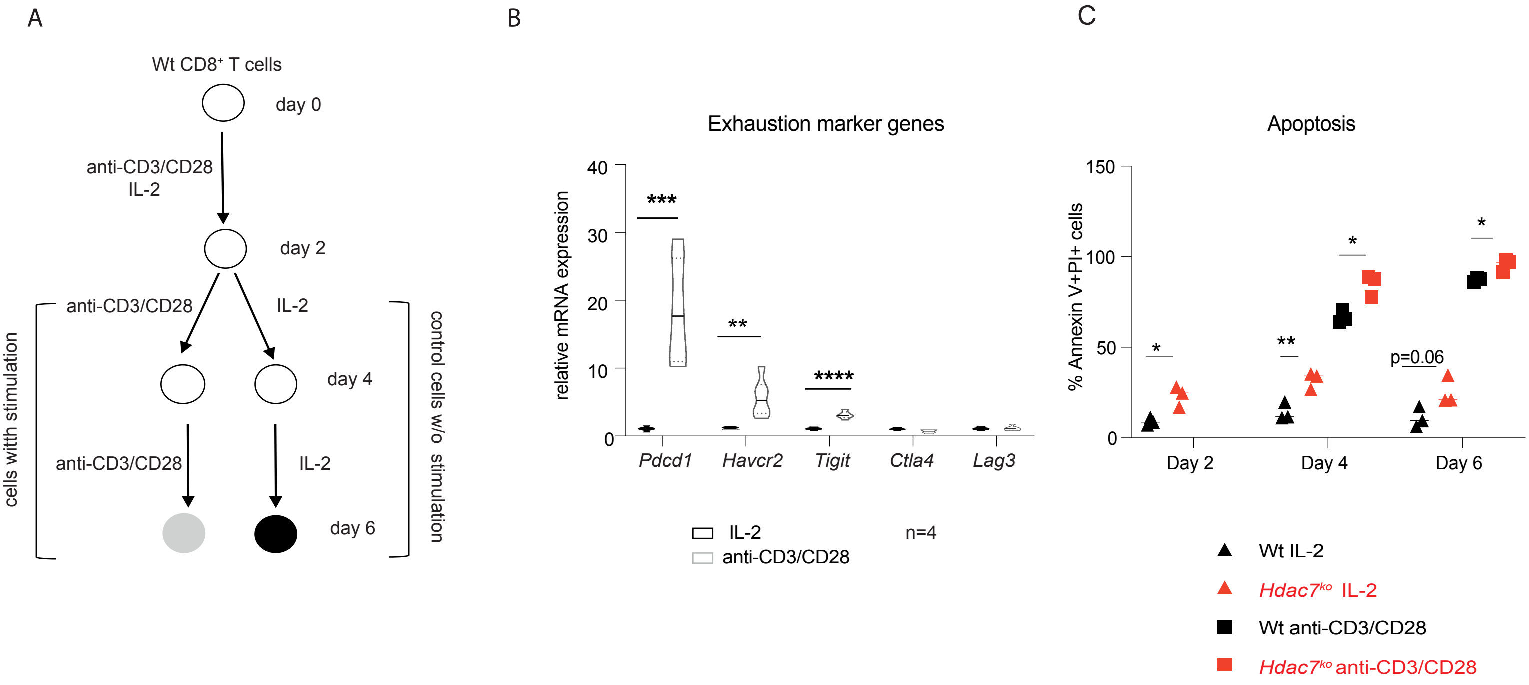


**Supplementary Figure 3: *Hdac7^ko^* CD8$^+$ T cells have increased apoptosis during chronic *in vitro* stimulation. (A)** Experimental set up for in vitro chronic stimulation model. Wt CD8$^+$ T cells were in vitro activated with anti-CD3/CD28 antibodies for 48 h and cultured in the presence of recombinant IL-2 only or restimulated with anti-CD3/CD28 antibodies for an additional 48 h. **(B)** Violin plots showing the expression of exhaustion marker genes *Pdcd1*, *Havcr2*, *Tigit*, *Ctla4* and *Lag3* in Wt CD8$^+$ T cells exposed to anti-CD3/CD28 stimulation or grown in the presence of IL-2 without repeated stimulation. Ct values were normalized to the Ct values of 36b4 and $2^{-\Delta Ct}$ was calculated. The expressions are shown relative to IL-2 controls. (n=4, multiple t test). **(C)** Dot plots showing the percentage of Annexin V$^+$PI$^+$ cells during repetitive *in vitro* stimulation of Wt and *Hdac7$^{ko}$* CD8$^+$ T cells. (n=3, multiple t test); *p<0.05, **p<0.01, ***p<0.001

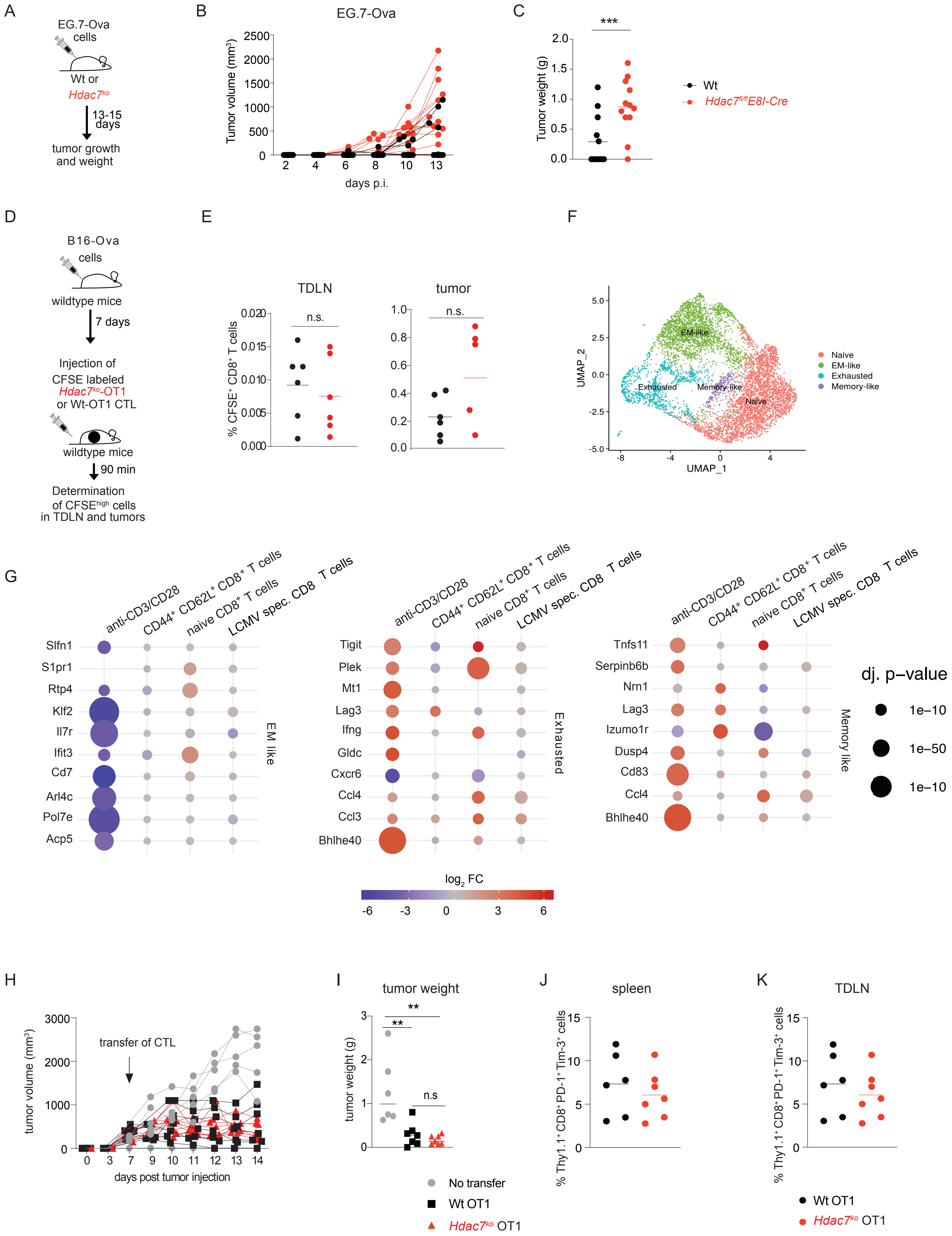
A
EG.7-Ova cells
Wt or
Hdac7ko
13-15 days
tumor growth and weight
B
EG.7-Ova
Tumor volume (mm3)
days p.i.
C
***
Tumor weight (g)
Wt
Hdac7fl/flE8I-Cre
D
B16-Ova cells
wildtype mice
7 days
Injection of CFSE labeled Hdac7ko-OT1 or Wt-OT1 CTL
wildtype mice
90 min
Determination of CFSEhigh cells in TDLN and tumors
E
TDLN
tumor
n.s.
% CFSE+ CD8+ T cells
F
UMAP_2
UMAP_1
EM-like
Exhausted
Memory-like
Naive
G
anti-CD3/CD28
CD44+ CD62L+ CD8+ T cells
naive CD8+ T cells
LCMV spec. CD8 T cells
Slfn1
S1pr1
Rtp4
Klf2
Il7r
Ifit3
Cd7
Arl4c
Pol7e
Acp5
EM like
Tigit
Plek
Mt1
Lag3
Ifng
Gldc
Cxcr6
Ccl4
Ccl3
Bhlhe40
Exhausted
Tnfs11
Serpinb6b
Nrn1
Lag3
Izumo1r
Dusp4
Cd83
Ccl4
Bhlhe40
Memory like
dj. p-value
1e−10
1e−50
1e−10
log2 FC
H
tumor volume (mm3)
transfer of CTL
days post tumor injection
I
tumor weight
**
**
n.s
tumor weight (g)
No transfer
Wt OT1
Hdac7ko OT1
J
spleen
% Thy1.1+ CD8+ PD-1+ Tim-3+ cells
K
TDLN
% Thy1.1+ CD8+ PD-1+ Tim-3+ cells
Wt OT1
Hdac7ko OT1

**Supplementary Figure 4: The anti-tumor immune responses of HDAC7 is CD8$^{+}$ T cell intrinsic. (A)** Wt and *Hdac7*$^{fl/fl}$*E8I-Cre* mice were intradermally (i.d.) injected with 1x10$^{6}$ EG.7-Ova cells. Tumor growth was followed for 14 days. **(B)** Tumor growth in Wt and *Hdac7*$^{fl/fl}$*E8ICre* mice (n=11-12). **(C)** Tumor weight in Wt and *Hdac7*$^{fl/fl}$*E8ICre* mice on day 14 after tumor inoculation (n=11-12 mice per group). **(D)** B16-Ova bearing Wt mice were i.v. injected with CFSE labeled Wt OT1 of *Hdac7*$^{ko}$ OT1 CTLs and sacrificed after 90 min. The frequency of CFSE-high cells was analyzed by flow cytometry. **(E)** The frequencies of CFSE-high CD8$^{+}$ T cells in tumor draining lymph nodes (TDLN) and B16-Ova tumors (n=5-6, multiple t test). **(F)** Unifold manifold approximation and projection (UMAP) plots displaying different subsets of B16-Ova infiltrating CD8$^{+}$ T cells[3]. **(G)** Top 10 regulated genes in anti-CD3/CD28 activated, CD44$^{+}$CD62L$^{+}$, naïve and LCMV-specific CD8$^{+}$ *Hdac7*$^{ko}$ CD8$^{+}$ T cells according to the gene sets obtained from B16-Ova infiltrating CD8$^{+}$ T cell subsets[3] (Wilcoxon test). **(H-K)** *Hdac7*$^{ko}$ mice were injected (i.d.) with 1x10$^{6}$ EG.7-Ova cells. 7 days after tumor inoculation, recipient mice were injected with 7x10$^{6}$ *in vitro* differentiated Wt OT1 or *Hdac7*$^{ko}$ OT1 CTLs. Tumor growth was followed for 7 days post CTL transfer. **(H)** Tumor growth and **(I)** Tumor weights in mice without transfer or transferred with Wt OT1 and *Hdac7*$^{ko}$ OT1 CTLs (n=6-7, multiple t test). **(J)** The frequency of transferred Thy1.1$^{+}$CD8$^{+}$PD-1$^{+}$Tim-3$^{+}$ Wt OT1 or *Hdac7*$^{ko}$ OT1 CTLs in spleen and **(K)** TDLN. (multiple t test $^{**}p<0.01$, $^{***}p<0.001$).

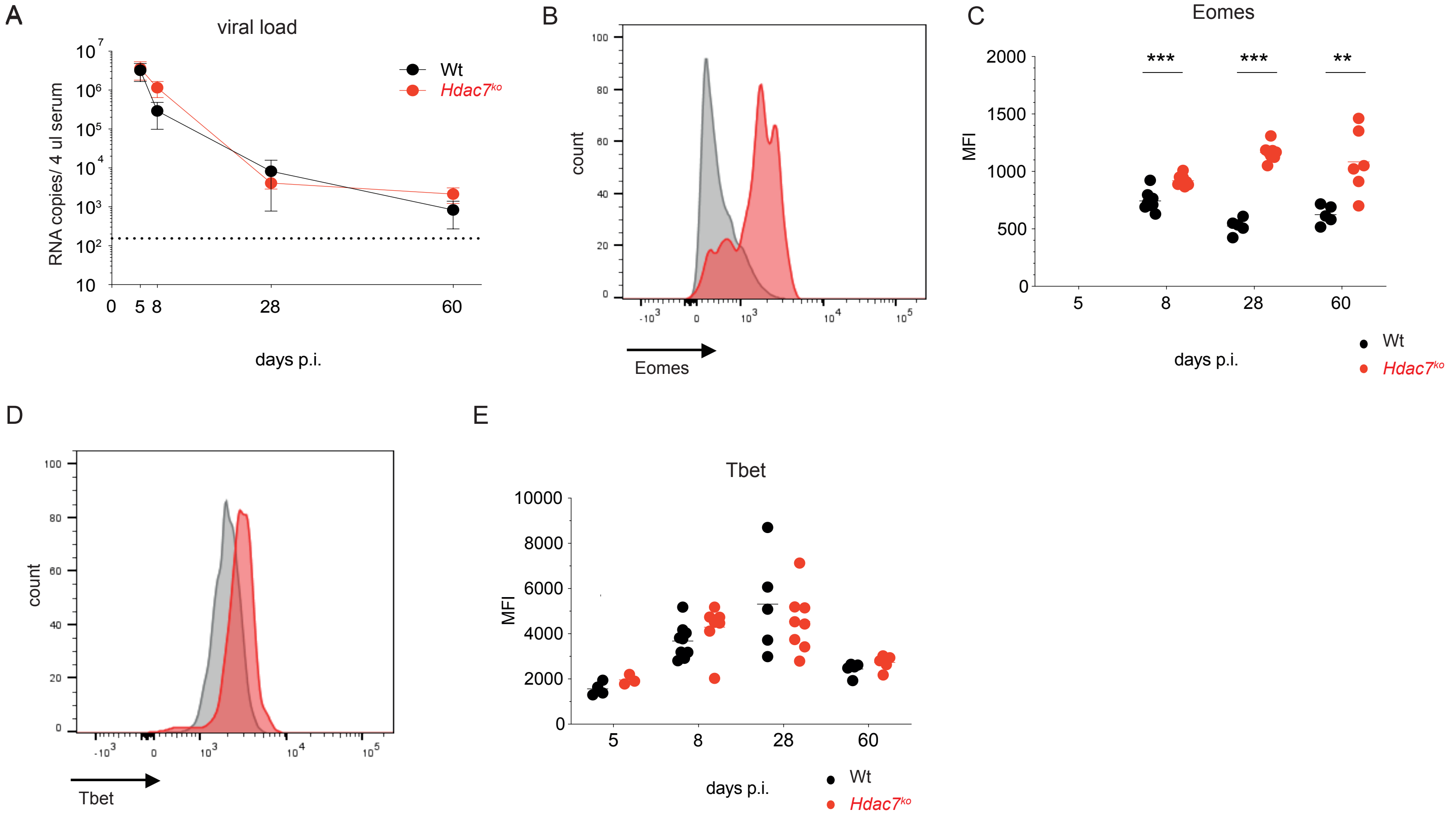


**Supplementary Figure 5: *Hdac7*$^{ko}$ CD8$^+$ T cells display increased Eomes expression during LCMV$_{Arm}$ infection. (A)** Viral load in the serum of LCMV$_{Arm}$ infected Wt and *Hdac7*$^{ko}$ mice as determined by RT PCR calculating LCMV$_{Arm}$ RNA copy numbers per 4 µl serum. **(B-E)** Representative histograms and dot plots displaying the mean fluorescence intensity (MFI) of **(B-C)** Eomes and **(D-E)** T-bet expression in CD3$^+$CD8$^+$Db-Gp33-streptamer$^+$ T cells isolated from the spleens of LCMV$_{Arm}$ infected Wt or *Hdac7*$^{ko}$ mice as assessed by flow cytometry (n=8-9, multiple t test); *p<0.05, **p<0.01, ***p<0.001.

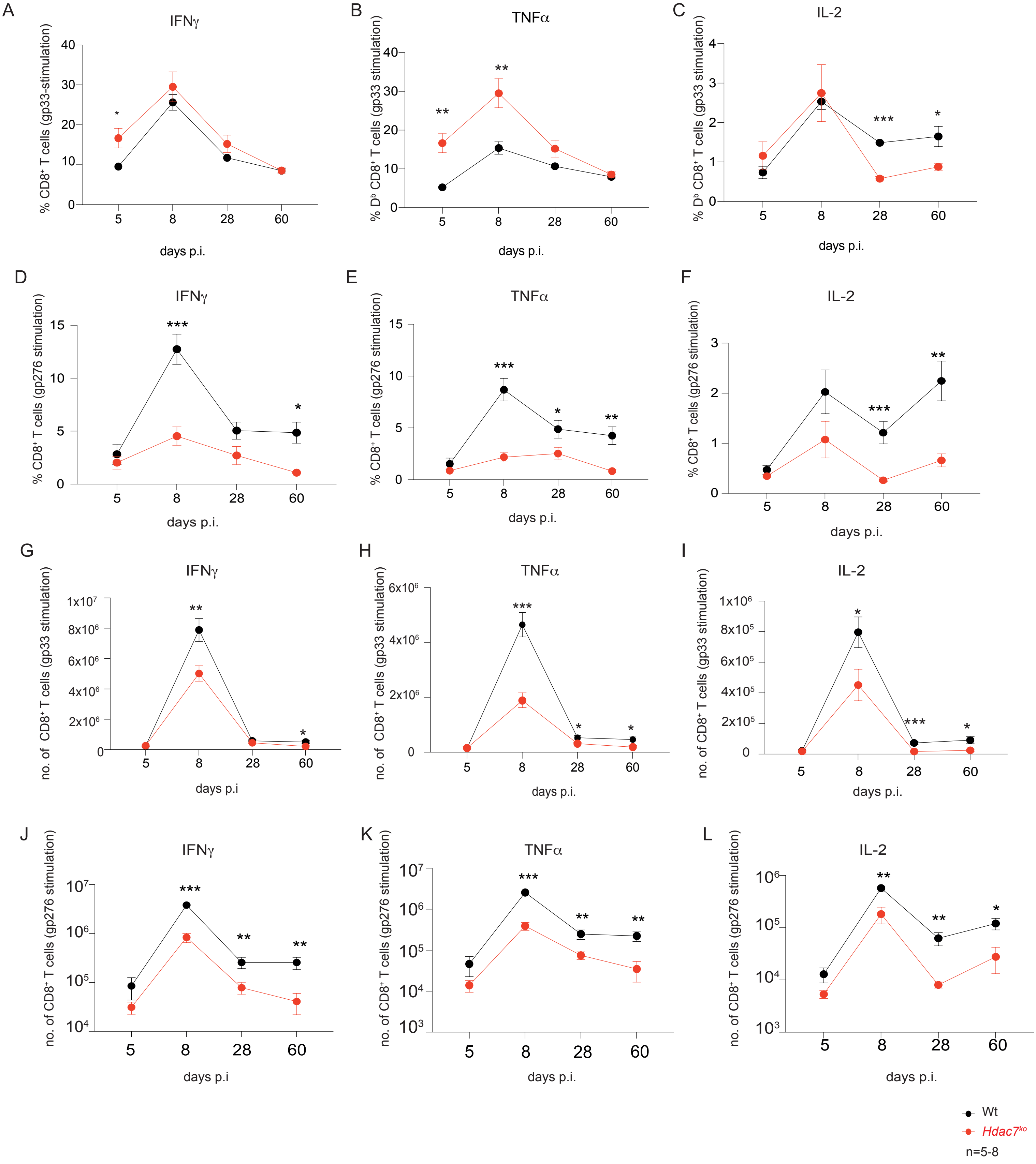


**Supplementary Figure 6: *Hdac7$^{ko}$* mice have altered frequencies of cytokine producing CD8$^+$ T cells during LCMV infection. (A-C)** The frequencies of IFNγ, TNFα and IL-2 producing Db-gp33 peptide stimulated Wt and *Hdac7$^{ko}$* CD8$^+$ T cells over the course of LCMV$_{Arm}$ infection. **(D-F)** The frequencies of IFNγ, TNFα and IL-2 producing gp276 peptide stimulated Wt and *Hdac7$^{ko}$* CD8$^+$ T cells over the course of LCMV$_{Arm}$ infection. **(G-I)** Cell counts of IFNγ, TNFα and IL-2 producing Db-gp33 peptide stimulated Wt and *Hdac7$^{ko}$* CD8$^+$ T cells over the course of LCMV$_{Arm}$ infection. **(J-L)** Cell counts of IFNγ, TNFα and IL-2 producing gp276 stimulated Wt and *Hdac7$^{ko}$* CD8$^+$ T cells over the course of LCMV$_{Arm}$ infection. (n=5-8, multiple t test) *$p<0.05$, **$p<0.01$, ***$p<0.001$.

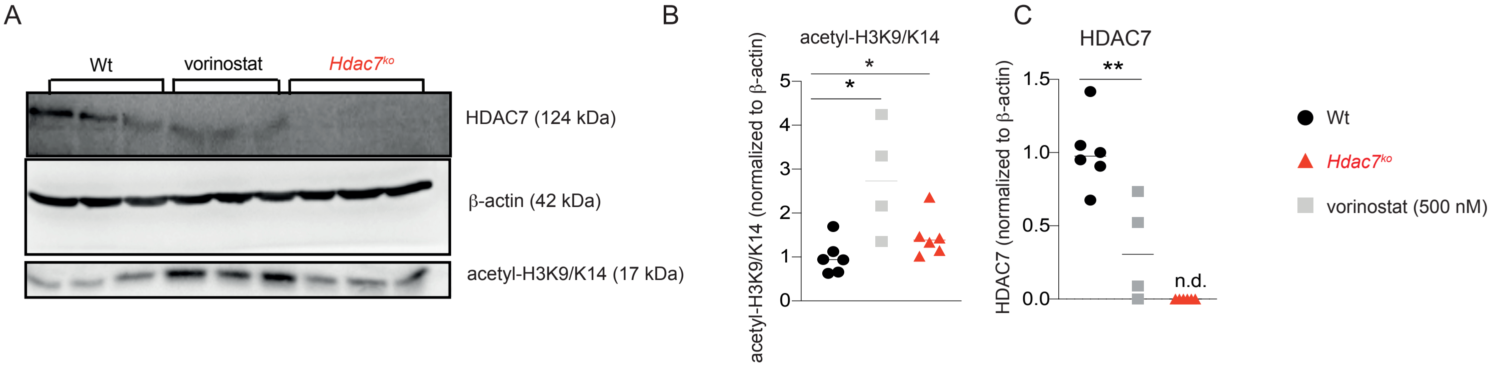


**Supplementary Figure 7: Vorinostat treatment of Wt CD8⁺ T cells results in reduced HDAC7 protein expression. (A)** Immunoblot analysis of HDAC7, acetylated-H3K9/K14 and β-actin in Wt CD8⁺ T cells activated with anti-CD3/CD28 antibodies and murine IL-2 for 48 h. Whole cell lysates were analyzed. Wt cells treated with vorinostat (500 nM) for 24 h served as positive control for acetylated-H3K9/K14 (representative of 3 independent experiments using biologically independent samples). **(B)** Fold-change protein expression of acetylated-H3K9/K14 and **(C)** HDAC7. Values were normalized to the intensity of β-actin bands (n=4-6, multiple t-test) *$p<0.05$, **$p<0.01$
